# LILAC: Enhanced actin imaging with an optogenetic Lifeact

**DOI:** 10.1101/2022.04.07.487416

**Authors:** Kourtney L. Kroll, Alexander R. French, Tobin R. Sosnick, Ronald S. Rock

## Abstract

We have designed an improved Lifeact variant that binds to actin under the control of light using the LOV2 protein. Light control enables one to subtract the pre-illumination signal of the unbound label, yielding an enhanced view of F-actin dynamics in cells. Furthermore, the tool eliminates actin network perturbations and cell sickness caused by Lifeact overexpression.

## Text

The actin cytoskeleton is central to a range of cellular processes, from cellular adhesion, shape and movement, to endo- and exocytosis, and cytokinesis. Dependable actin visualization is vital for various fields of biological research^1^. One label, Lifeact, has become the gold standard for labeling actin filaments in cells, being cited in over 1800 papers^2^.

However, recent work has shown that Lifeact has unintended effects, such as Drosophila sterility, mesenchymal stem cell morphology changes, and altered actin patch and cytokinetic ring dynamics in fission yeast^3-5^. These side effects are likely due to competitive binding between Lifeact and other important proteins, such as cofilin and myosin, which also bind actin filaments at the D-loop^6,7^. To address this serious problem and develop a new optogenetic tool, we designed LILAC (light-induced localization to actin) that has low actin affinity in the dark. Upon excitation, the photo-activated Lifeact diffuses within a second from the cytosol to actin filaments, labeling them for imaging. LILAC photocycles and returns to the low affinity state after a minute which minimizes side effects. In addition, the light-controlled aspect greatly improves the imaging capability.

In our design, we employed LOV2, the second light-sensitive light-oxygen-voltage (LOV) domain from phototropinI in oats which is involved in the phototropism response in plants. In response to blue light, the flavin chromophore forms an adduct with C450, resulting in conformational changes within the core^8-9^. This change disrupts the seven residue N-terminal A’ɑ helix and 24 residue C-terminal Jɑ helix, causing them to unfold^10^. This light-dependent conformational change of LOV2 has been harnessed to engineer a variety of optogenetics tools^17-21^.

The dark state recovery of LOV2 has a time constant of 81 sec for the wild type protein at room temperature and pH 8^11^. This time constant is tunable over several orders of magnitude through mutations and buffer conditions^12,13^. Additionally, mutations to mimic constitutive light-exposed, unfolded (I539E) and the dark, folded state (C450A) are well characterized^14-16^.

Using previous LOV2-based optogenetic platforms as a guide and minimizing disruption of the hydrophobic contacts between the Jɑ helix and the core of LOV2, we designed LILAC (Figure 1a). This design is intended to sterically prevent binding to actin by caging Lifeact, when the Jɑ helix is folded (Fig. 1b, red). This caging is achieved by positioning the hydrophobic residues of Lifeact, which are necessary for actin binding in cells^6,7^, towards the core of LOV2. Upon unfolding of the Ja helix, Lifeact is uncaged and able to bind actin filaments.

**Figure 1:**
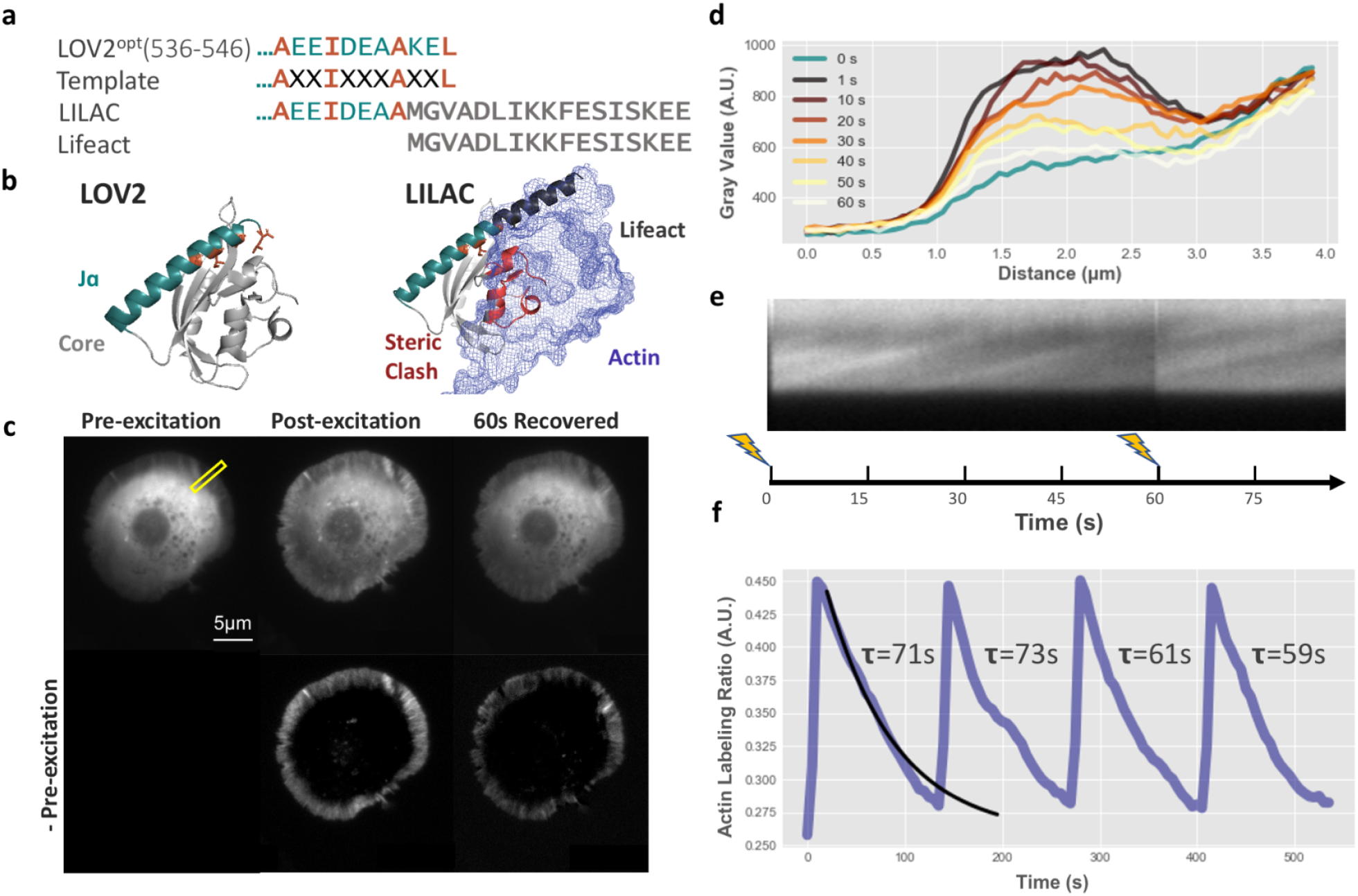
LILAC reversibly binds actin in S2 cells. **a**, Sequences of the LOV2 C-terminal Jɑ helix and Lifeact. The template was created by keeping hydrophobic residues (blue) which interact with the core of LOV2, and Lifeact was integrated to minimize disrupting those Interactions. Grey:LOV2, turquoise: Jɑ helix, orange: key hydrophobic residues, black: Lifeact. **b**, Structural representations of LOV2 (left) and LILAC (right). Actin (blue) is shown in bound to Lifeact from PDB 7AD9. LOV2 residues that sterically clash with actin in the dark state are shown in dark red. **c**, TIRF images of an S2 cell that is expressing LILAC taken before, during, and after excitation with blue light. **d**, Line scan of cell area highlighted in yellow in **c** before blue light excitation at 0 seconds and 1 to 60 seconds post excitation, where 0 μm is outside of the cell. Scale bar is 5 μm. **e**, Kymograph of the outer cell edge highlighted in yellow in (**c**), where the top is the inside of the cell, and the bottom is outside the cell. **f**, Actin labeling ratio of a repeatedly excited cell.

We used TIRF imaging to test filament labeling in the light and dark. After blue light excitation, *D. melanogaster* S2 cells have increased fluorescence in the outer lamellipodial actin ring as compared to the dark state (Figure 1c, Supp. Movies 1,2). This increased lamellipodial fluorescence reverses with a time constant similar to WT LOV2, suggesting that the reversion of the LOV2 domain to its dark state drives unbinding from the actin (Figure 1d). Additionally, by subtracting the pre-excitation image, we effectively eliminate the background fluorescence from the cytosolic, unbound LILAC. This subtraction yields a substantially improved image that highlights the structures of interest (Figure S1a-b), and reduces the influence of variations in cellular thickness. Subtraction of the pre-excitation image also enhances the appearance of retrograde flow in kymographs (Figure S1c-d). These enhanced Kymographs show retrograde flow of actin at a rate of 3.5 ± 0.12 μm/min, which is similar to the rate of 4.0 ± 0.44 found by Rogers et. al. (Figure 1e).

The LILAC labeling of the lamellipodium decays over time, implying reversible, light-activated labeling. To quantify these images, we use the actin labeling ratio (ALR), defined as the ratio of average fluorescence in the outer ring, divided by the average fluorescence of the inside of the cells over time (Figure 1f, S2). This quantity sharply increases immediately after excitation, followed by the recovery back to nearly pre-excitation levels. We define the switching ratio as the ratio of the actin labeling intensity post and pre-excitation. This photo-excitation can be repeated multiple times, with no reduction in efficacy (Figure 1f).

Individual S2 cells express varying amounts of the LILAC. As the overall cell intensity increases, the relative actin labeling intensity decreases (R^2^=0.27, Figure S3a). This decrease is likely due to an increase in the excess cytosolic LILAC with increased expression. We also find that the switching ratio of actin labeling in the light versus dark has weak correlation with cell intensity (R^2^=0.13, Figure S3b). This switching ratio relationship suggests that at high expression, there is a high background from unbound monomer; however, when comparing the dark and light states, one can still readily identify the actin-rich areas of the cell. Additionally, we found that the maximum actin-labeling ratios of LILAC are similar to Lifeact, implying that the activated LILAC binds F-actin with a similar affinity as Lifeact (Figure S3c). LILAC has a dark vs. light switching ratio of ∼1.6, while expectedly, Lifeact and LILACI539E exhibit no change in the light with switching ratios of ∼1 (Figure S3d).

We next examined the dark state recovery process to determine how rapidly the cell will reset after imaging. From actin labeling ratio traces, we determine the recovery time constant associated with each cell. On average, cells recover with a time constant of 63 seconds, close to the characteristic 80 second *in vitro* time constant of LOV2^11^ (Figure S4). We repeated this measurement with our T406A/T407A mutation, which stabilizes the N-terminal A’ɑ helix, reduces the recovery time constant, and can increase caging^11,17^. The average time constant decreased to 50 seconds, similar to the 54 second time constant measured previously *in vitro*^11^. Unexpectedly, we observed a considerable amount of cell-to-cell variance in the recovery time, but for a given cell, the first and second recovery times were strongly correlated (R^2^=0.70, Figures 1f, S4). This correlation suggests that each cell has an intrinsic time constant that is linked to cellular state (e.g., metabolite concentrations or pH), given that LOV2 dark state recovery is sensitive to buffer conditions^23^. To determine whether we could easily perturb these recovery times, we added 1 mM imidazole to cell media. Imidazole treatment greatly reduced the recovery time constants across all cells. Imidazole can therefore be used to enhance dark-state recovery as desired.

Next, we examined the effects of high mCherry-Lifeact expression levels on the morphology of insect S2 cells using total internal reflection fluorescence microscopy (TIRF). We found that at low expression levels, cells are round with the characteristic lamellipodial actin ring at the outer edge. At increasing concentrations of mCherry-Lifeact, however, cells begin to form long F-actin bundles and protrusions (Figure 2a). Such cells have an irregular shape and have impaired spreading onto the coverslips. This enhanced F-actin assembly could be due to the competition between Lifeact and cofilin^6^, which in turn reduces F-actin turnover.

**Figure 2:**
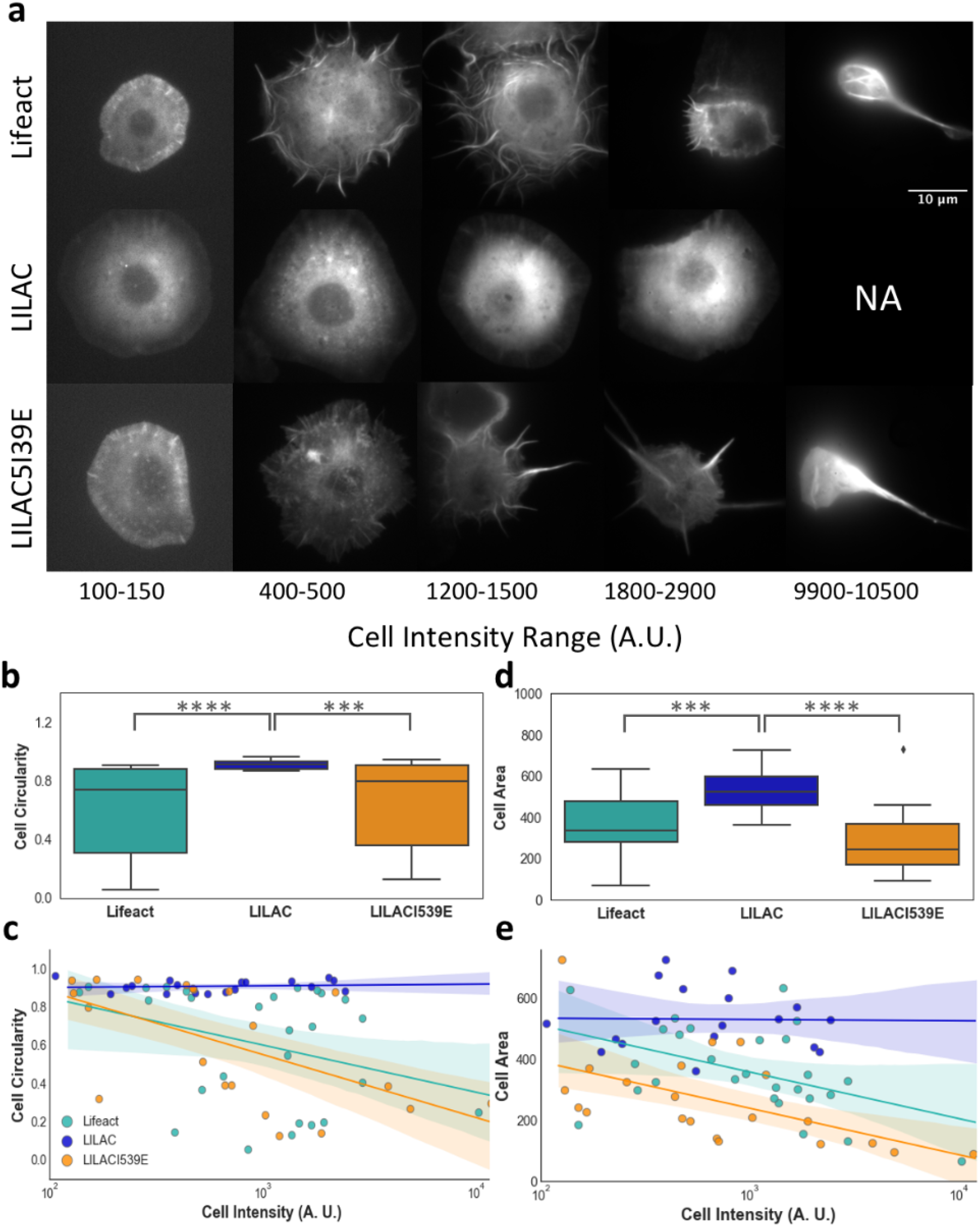
LILAC reduces the concentration dependent side effects of Lifeact. **a**, Cells at various Intensities expressing Lifeact, LILAC, and LILACI539E. Cell circularity (**b**, P = 1.57*10^−4^) and cell area **(e**, P = 3.77*10^−6^) of Lifeact, LILAC, LILACI539E. The center line represents the median. Boxes span the 25th and 75th percentiles. Whiskers extend from the 25th percentile - 1.5x the interquartile range (IQR) and From the 75th percentile + 1.5x IQR. Data beyond the whiskers represent outliers and are plotted individually. Cell circularity **(c)**, cell area **(e)** as a function of intensity with a linear regression fit to log-transformed data and shaded regions indicating the 95% confidence interval. For **b**, and **d**, P-values were determined initially using a one-way ANOVA test (*n*=27 for Lifeact, *n*=18 for LILAC, *n*=23 for LILACI539E). A post-hoc Dunn test was used to determine pairwise p-values. (***=P≤0.001,* ***=P≤0.0001)

LILAC greatly reduces these morphological effects induced by Lifeact expression. At the highest level of expression, LILAC expressing cells maintain their round shape and lack the prominent F-actin bundles (Figure 2a). In addition to the circularity being significantly reduced with Lifeact in comparison to LILAC (P = 1.57*10^−4^, Figures 2b,c), we also observe that circularity weakly depends on LifeAct expression level (R^2^ = 0.12), but does not depend on LILAC expression level (R^2^ = 0.012). Lifeact expressing cells also have reduced spread area as a function of expression level (R^2^ = 0.21). This effect is absent when expressing LILAC (R^2^ = 2.8*10^−4^, Figures 2d,e). These observations quantitatively suggest that Lifeact alters the actin cytoskeleton and the resulting cell morphology whereas LILAC does not.

To test whether the lack of cytoskeletal artefacts in LILAC-labeled cells was due to its ability to effectively cage the Lifeact peptide, we evaluated circularity and cell area in a LILAC containing an I539E mutation in the LOV2 domain. This mutation forces the unfolding of the Jα helix, rendering the LILACI539E constitutively active^14-16,18^. As expected, LILACI539E recapitulates the morphological defects seen in S2 cells overexpressing mCherry-Lifeact (Figure 2a). Similarly to mCherry-Lifeact, circularity (R^2^ = 0.40, Figures 2b,c) and cell area (R^2^ = 0.36, Figures 2d,e) correlate with expression level. Overall, our results show that LILAC is nearly inert in the dark, even when highly overexpressed, and that this is likely due to effective caging of the Lifeact peptide.

In summary, LILAC has great potential utility as a new tool for actin imaging. By subtracting the pre-excitation image, we eliminate the cytosolic background and obtain superior images of the actin cytoskeleton alone. Recovery times can be tuned by adjusting solvent conditions, allowing for slow recovery for recording movies of actin dynamics, or rapid recovery when minimal perturbation is required. Moreover, LILAC eliminates Lifeact induced side-effects when cells are grown in the dark. In the future, we envision that a spatiotemporally controlled actin binding protein can be used to control actin cross-linking and patterning, and the subcellular recruitment of actin binding proteins.

## Acknowledgements

This work was supported by grant numbers EB009412, GM055694 and GM124272 awarded by the National Institutes of Health, as well as the David Grier Prize for Innovative Research in Biophysical Sciences.

## Author Contributions

Author contributions: K.L.K., A.R.F., T.R.S., and R.S.R. designed research; K.L.K. performed research; K.L.K., T.R.S., and R.S.R. contributed new reagents/analytic tools; K.L.K., T.R.S., and R.S.R. analyzed data; and K.L.K., A.R.F., T.R.S., and R.S.R. wrote the paper.

## Methods

### DNA preparation

A LILAC G-block was purchased from Integrated Dna Technologies and incorporated into a pet21 vector containing 6xHis-mCherry-TEV using Gibson Assembly containing AsLOV2(401-543) with the mutations G528A/I532A/N538E for improved switching^1^ to make LOV2^opt^. Next, we used gibson assembly to incorporate mCherry-TEV-LILAC into a pMT vector from the Glotzer laboratory (UChicago). The TEV sequence was then replaced with a GS linker using the NEB Q5 protocol. The T406A/T407A and I539E mutations were created using the NEB Q5 protocol.

### Cell maintenance

Insect S2 cells from the Fehon Lab (UChicago) were maintained in Schneider’s insect media (Sigma) with insect media supplement (Sigma) and 50 units/mL penicillin and 50 μg/ml streptomycin (Thermo). Cells were split 1:6 every 4 to 6 days at around 10-15 million cells/ml at 20 ml in T75 flasks.

### Transfection and Imaging Preparation

Cells were transfected as previously described^2^. In short, 2 ml of fast-growing cells at 1.35 million/ml to a 6 well plate. DDAB at 250 μg/ml was added to the media in a 1:2 ratio and left for ten minutes. Then, the DDAB-media mixture was added to 500ng of pMT vector to reach 150 μl and left for five minutes. This was added dropwise to the cell. One day later, cells were induced with 1.5 μl of 0.7M CuS04. Cells were transferred one day later on glass-bottom dishes coated with 15 μl of 0.5 mg/ml concanavalin A^3^. For Figure S3, 1M sterile filtered imidazole was added to media to a final concentration of 1mM. After 30 minutes to allow for cell spreading, cells were imaged.

### Imaging

Dishes were imaged using an ×100, 1.65 N.A. objective (Olympus) on a custom-built total internal reflection microscope using an electron-multiplying charge-coupled device (EMCCD) camera (iXon; Andor Technologies) with a mCherry specific emission filter. This microscope was controlled with the open source Micro-Manager program (https://micro-manager.org/). Cells excited twice were excited first with 561 nm, then with 488nm and 561nm, and recovery images were taken with just 561nm excitation, all at 1% power for 500 ms exposure with an EM gain of 200 with no delay time between images. Cells excited more than twice were imaged in exactly the same manner with 5 s between images instead of 500 ms. Cells in Figure 2 were imaged on the same microscope on the same day to best guarantee cells of the same brightness are expressing similar protein levels.

### Image Analysis

Outer and inner rings of the cells were traced by hand and kymographs were created in Fiji (https://imagej.net). Brightness, cell perimeter, area, circularity, and roundness values were exported from Fiji for inner and outer cell areas for each cell. Time constants were calculated by fitting actin labeling ratio to a single exponential. Data analysis and figures were made using Python scripts.

We quantified morphological changes by calculating the cell circularity, area, and roundness. When cells are circular, this value is 1, and when cells have large perimeters, it approaches 0. R^2^ values were calculated using the logarithm of the cell intensity and the circularity/cell area.

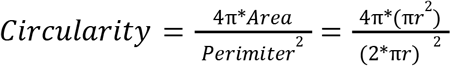

## Data Availability

The datasets and code generated during the current study are available from the corresponding authors on reasonable request.

---

**Figure S1:**
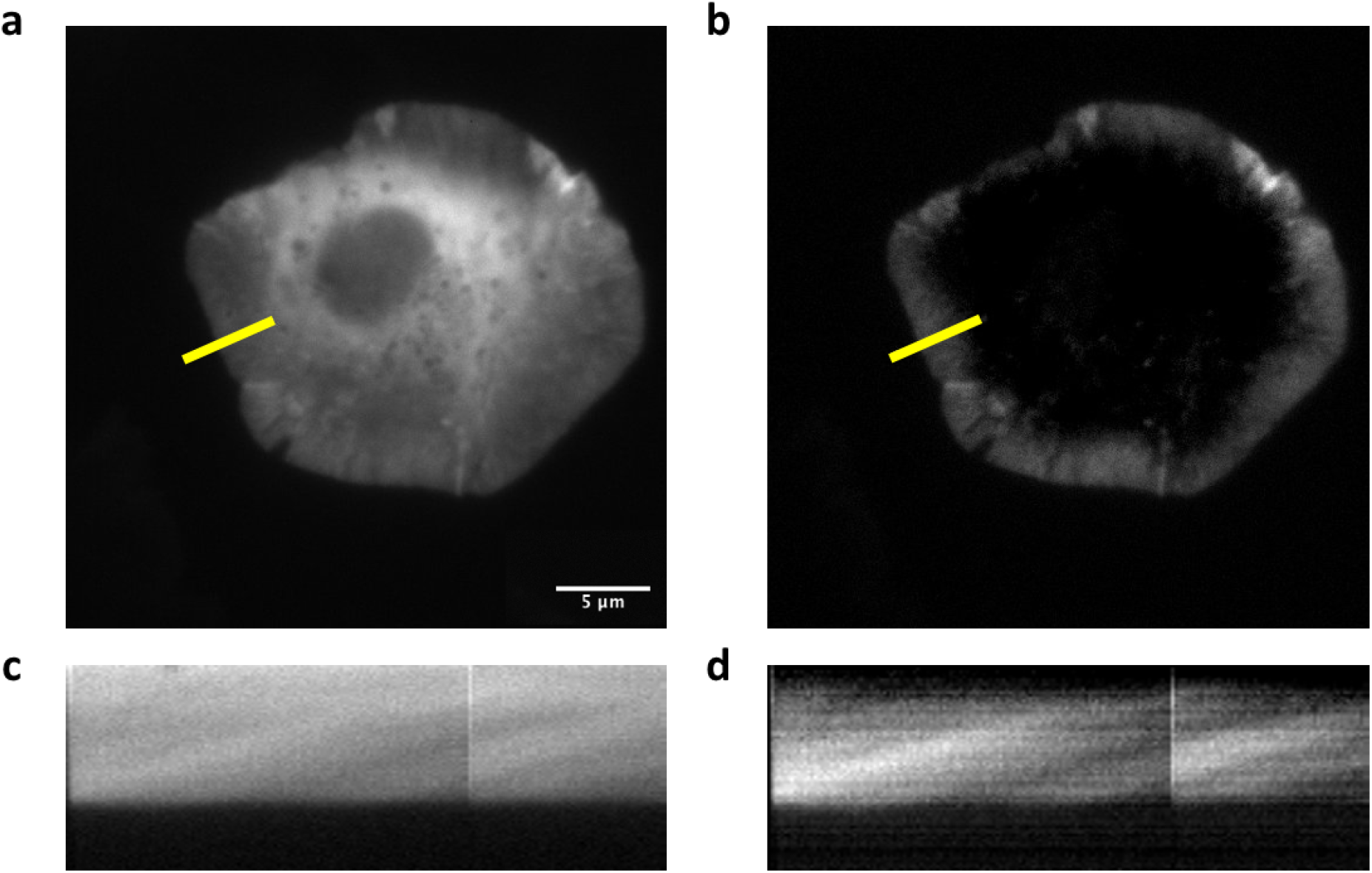
LILAC highlights actin structures through background subtraction. TIRF image of an S2 cell expressing LILAC without **(a)** and with **(c)** pre-excitation image background subtraction to eliminate cytosolic background. Kymographs of the region in yellow without **(c)** and with **(d)** pre-excitation background subtraction

**Figure S2:**
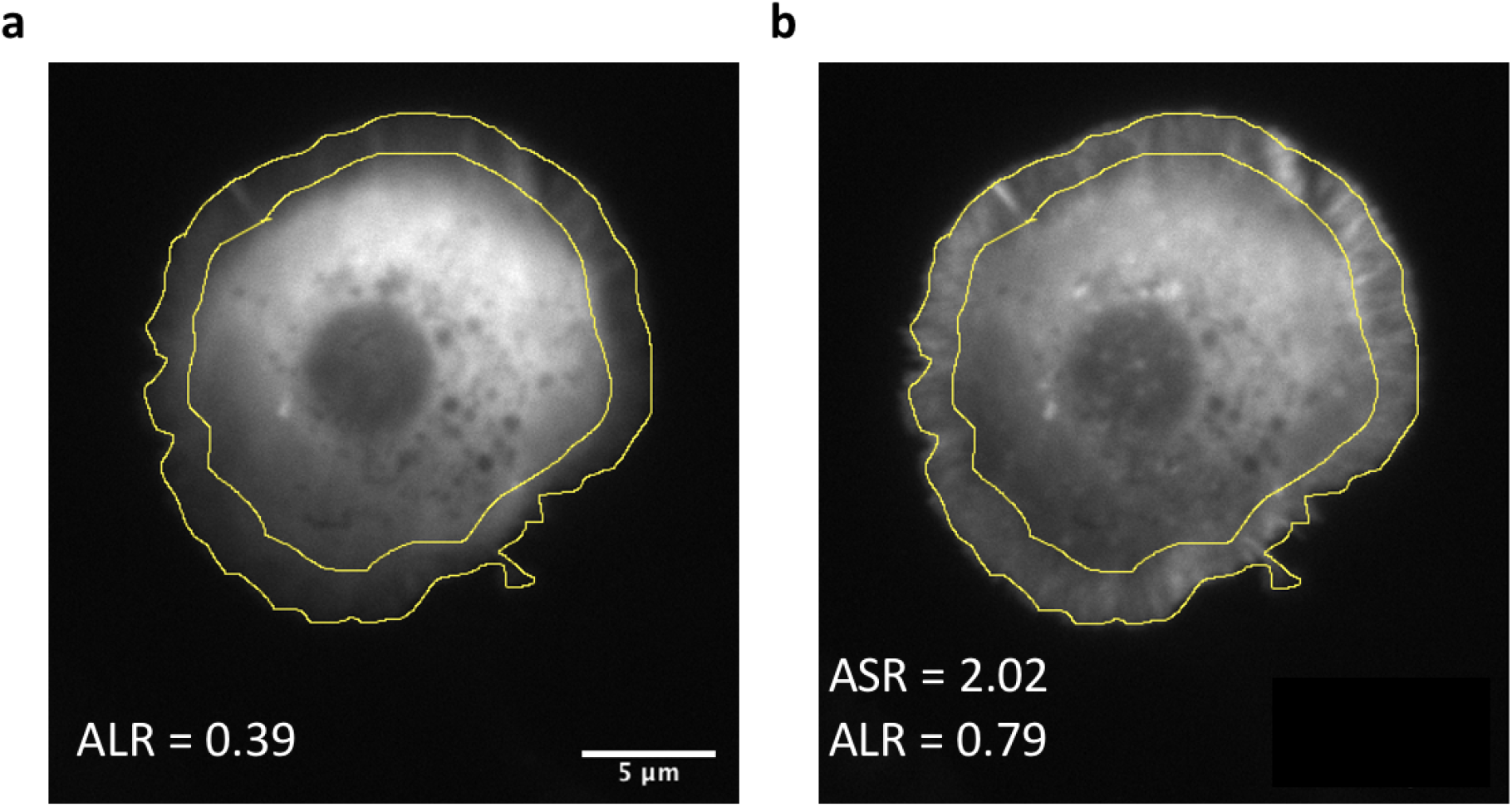
Image processing of LILAC cells to obtain labeling and switching ratios. An S2 cell expressing mCherry-LILAC before **(a)** and after **(c)** blue light excitation. The outer and inner rings of cells are hand traced in FIJI (yellow). Actin labeling ratio: ALR = lntensity_inner_/lntensity_outer_, actin switching ratio: ASR = ALR_post_/ALR_pre_.

**Figure S3:**
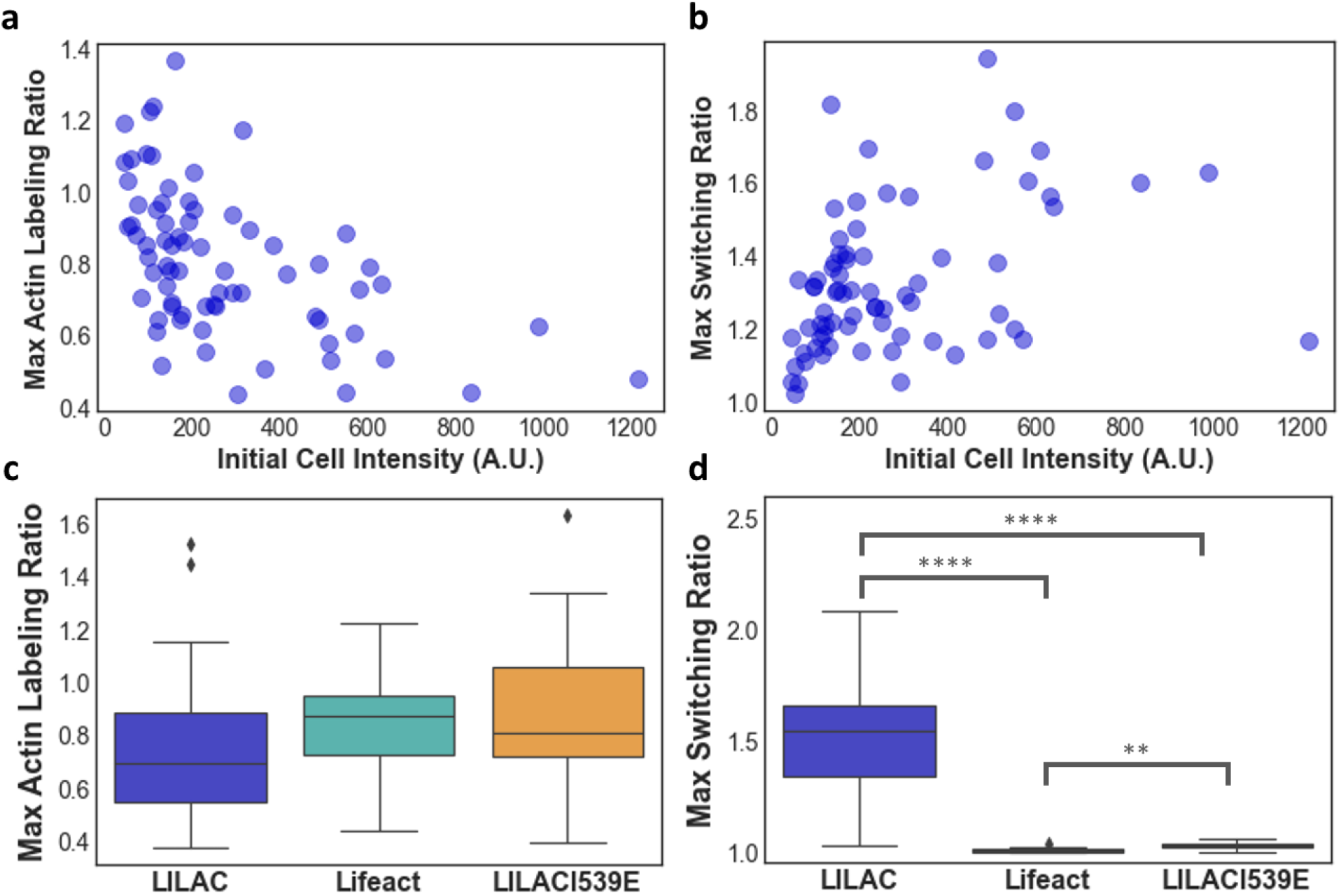
Quantification of labeling and switching of LILAC. Max actin labeling ratio (**a**, R^2^ = 0.27) and switching ratio (**b**, R^2^=0.13) of LILAC expressing cells as a function of cell intensity. Maximum actin labeling (**c**, P = 0.11) and switching ratios (**d**, P = 9.66* 10^-9) of LILAC, Lifeact, and and the light-mimic mutant, LILACI539E. The center line represents the median. Boxes span the 25th and 75th percentiles. Whiskers extend from the 25th percentile - 1.5x the interquartile range (IQR) and from the 75th percentile + 1.5x IQR. Data beyond the whiskers represent outliers and are plotted individually. For **c** and **d**, P-values were determined initially using a one-way ANOVA test (*n*=27 for Lifeact, *n*=18 for LILAC, *n*=23 for LILACI539E). A post-hoc Dunn test was used to determine pairwise p-values. (**=P≤0.01, ***=P ≤0.001, ****=P ≤0.0001)

**Figure S4:**
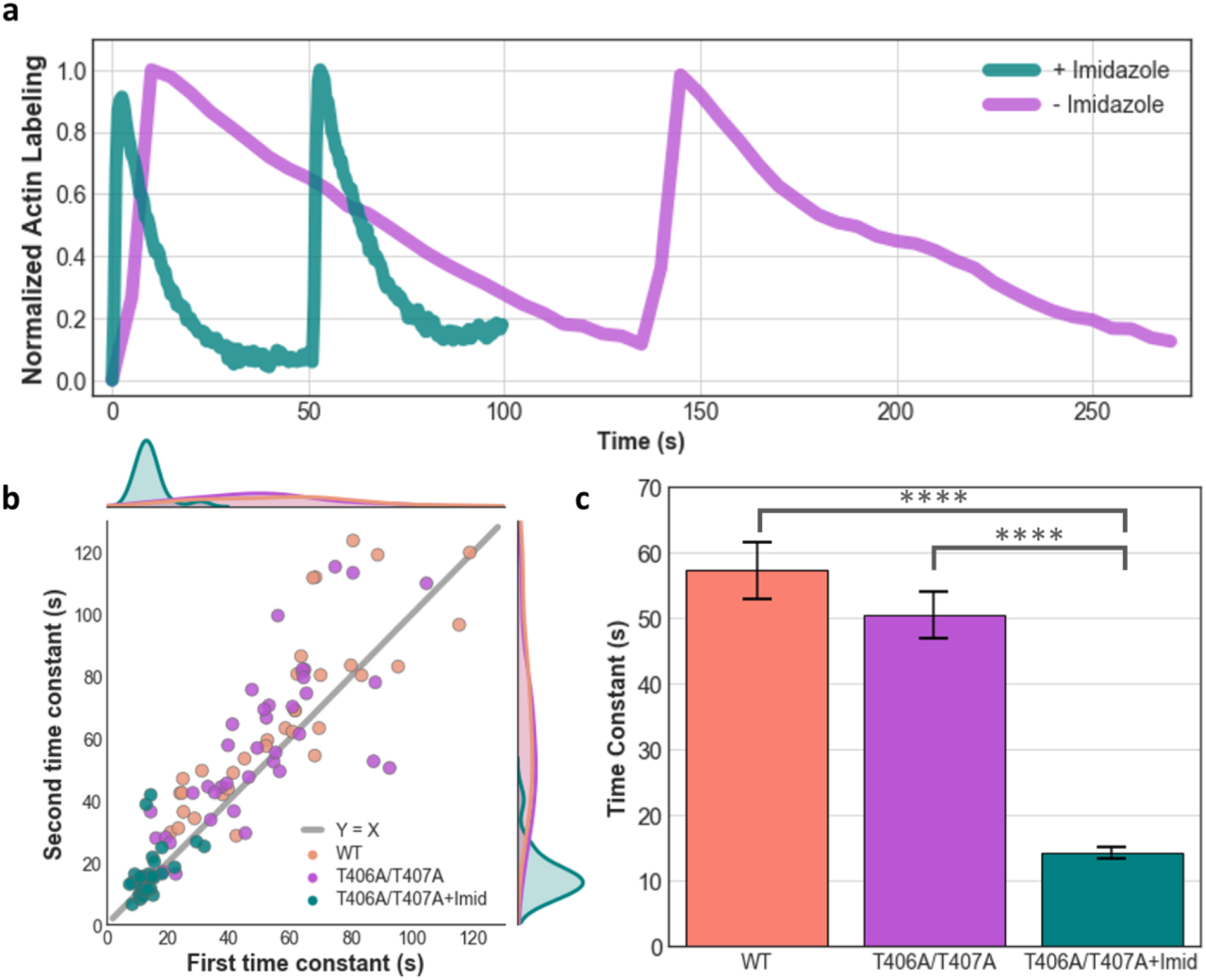
Dark state recovery time constants are cell specific and tunable. **a**, Min-max normalized actin labeling ratio traces of LILACT406A/T407A expressing cells with and without 1mM imidazole. Cells were activated with blue light every 60 and 130 seconds respectively. **b**, First and second recovery time constants for individual cells expressing WT LILAC (R^2^=0.70), LILACT406A/T407A (R^2^=056), and LILACT406A/T407A with 1mM imidazole (R^2^=0.19). Marginal kernel density estimates plotted along edges. **d**, Barplots of time constants of LILAC (*n*=34), LILACT406A/T407A (*n*=38), and LILACT406A/T407A with 1mM imidazole (*n*=33) with error bars representing standard error (****=P≤0.0001).

**Movie S1: Light-induced labeling of lamellopial with LILAC**. TIRF imaging of an S2 cells, expressing mCherry tagged LILAC. Blue light (488nm) is pulsed for 0.5 seconds after the first fame, and again at one minute.

**Movie S2: Labeling with LILAC allows for enhanced imaging with pre-excitation subtraction**. TIRF imaging of an S2 cells expressing mCherry tagged LILAC., with the pre-excitation image subtracted from all images in the movie. Negative values are all floored to 0 (black). Blue light (488nm) is pulsed for 0.5 seconds after the first fame, and again at one minute.

